# Long-term temperature trend in Kamchatka supports expansion of harmful algae

**DOI:** 10.1101/2022.03.24.485652

**Authors:** Kanat Samarkhanov, Yersultan Mirasbekov, Ayagoz Meirkhanova, Adina Zhumakhanova, Dmitry Malashenkov, Alexander Kovaldji, Natasha S. Barteneva

**Author notes:** Co-Senior authors Kanat Samarkhanov, Natalie (Natasha) S. Barteneva. These authors contributed equally to this study (Kanat Samarkhanov, Yersultan Mirasbekov). IMIM, Heidelberg University, 69117, Heidelberg, Germany.

## Abstract

Ocean coastal ecosystems are changing, and global shifts in temperature lead to the expansion and intensification of harmful algae. In conjunction with anthropogenic effects it may result in future exacerbation of harmful algal blooms. Here we use the 2002-2020 years record of surface ocean temperature data retrieved from Sentinel-2 satellite to examine the recent temperature trend in Avacha Bay, Kamchatka Peninsula. Satellite analysis demonstrated a temperature increase trend in ocean surface water during spring and summer months and detected algal bloom in July 2020 preceding a mass death of marine benthic life in September-October 2020. Using 16S rRNA and 18S rRNA gene amplicon nanopore-based sequencing, we analyzed microbial and microalgal communities in the water samples from area of 2020 algal blooms. Our results suggest the presence of potentially toxic and bloom-forming algae from genera related to former HABs (harmful algal blooms) in the Avacha Bay region. A better understanding of the potentially toxic algae phytoplankton composition in the shifting temperature environment and time-series monitoring of HABs is of utmost importance for scientific community. We suggest that satellite analysis in combination with eDNA monitoring by nanopore-based sequencing represents promising option to detect potentially toxic algae and follow bloom development.

## Introduction

The harmful algae expand over their spatial range limits, harmful algal blooms (HABs) happen in different coastal areas, and the underlying causes and factors influencing the dynamics of toxic algae are targets of many studies (Hallegraeff, 2010; Bucci et al., 2020; Hays et al., 2005; Gobler et al., 2017; Ho et al., 2019; Griffith et al., 2020; Trainer et al., 2020; Errera et al., 2014). The temperature of the upper ocean is a fundamental feature controlling phytoplankton metabolic processes (Raven, and Geider, 1988; Moisan et al., 2002; Thomas et al., 2012) and a major metric that could be used for predicting HABs (Ralston et al., 2014). The near-surface layers of the ocean respond more rapidly and strongly to warming and are more sensitive to freshwater inputs that influence salinity and stratification (Errera et al., 2014). Moreover, ecosystem-level responses to temperature change can disrupt the seasonal timing of recurring processes (Portner, and Farrell, 2008) and lead to a generation of long-lasting seasonal windows for blooms to occur (Hallegraeff, 2010). Widespread blooms of harmful marine dinoflagellates (*Karenia* and *Alexandrium* spp.) and diatoms (*Pseudo-nitzschia* spp.) (Kurenkov, 1974; Konovalova, 1992;1993; 2000; Konovalova, and Sizykh, 2020; Orlova et al., 2002; 2010; Selina et al., 2006; Sakamoto et al., 2021), and diazotrophs such as *Anabaena* spp. (Konovalova, 2000; Walsh et al., 2011) were described during recent years in the coastal areas of Kamchatka and Avacha Bay.

HABs lead to shellfish poisoning, fish kill, and economic damage (Landsberg, 2002). There are >200 species of marine potentially toxic microalgae (∼4% of the ∼5000 phytoplankton species) (Landsberg, 2002; Landsberg et al., 2014; Hallegraeff et al., 2021), and this number is continually increasing to include algae previously considered benign (James et al., 2003; Dai et al., 2019). HABs often have a mixed origin. A better understanding of the potentially toxic algae phytoplankton composition in temperature shifting environment is of utmost importance for the scientific community. Traditional identification methods, including light microscopy, provide limiting capabilities for the detection of mixed HABs. Many dinoflagellate species cannot be unambiguously separated under conventional light microscopy. Thus, *Alexandrium* spp. share many morphological features with other thecate dinoflagellates making identifying microalgae responsible for the specific blooming event on the basis of morphological traits and only light microscopy impossible (Kim et al., 2017). The cysts of some dinoflagellate species such as *Karenia mikimotoi* (Liu et al., 2020), *Akashiwo sanguinea* (Tang and Gobler, 2015), *Cochlodinium polycricoides* (Liu et al., 2020; Tang and Gobler, 2012) lack distinctive morphological traits and cannot be accurately identified using light, epifluorescence and even electron microscopy, which are costly and time-consuming. As a result, there has been confusion as to how many *Karenia* species were present during PSP (paralytic shellfish poisoning) events (Chang et al., 2004). Molecular identification of toxic HAB species has become important in the last few decades as it offers a faster cost-effective alternative (Adachi et al., 1996; Godhe et al., 2006; Gray et al., 2003; Galluzzi et al., 2004; Casper et al., 2004; Dyhrman et al., 2006; Kamikawa et al., 2008; Carneau et al., 2011; Anderson et al., 2012; Jedlicki et al., 2012; Lee et al., 2020). Though PCR-based technologies are now widely used in research, their role in HAB monitoring would require parallel multiplexing of numerous assays making this approach impractical (Bott et al., 2010; Hatfield et al., 2020). The PCR-based methods are target-specific, may introduce amplification bias, and may not amplify driver of algal bloom (Krehenwinkel et al., 2017). On the other hand, the recent development of next-generation sequencing (NGS) technologies led to their availability for field analysis and offered an enormous perspective for HAB monitoring (Jung et al., 2018; Hatfield et al., 2020). Ocean color satellite images is another key to studying the temperature trends and response of marine phytoplankton to climate variability and change (Groom et al., 2019), plankton distribution in ocean, and detecting and predicting HABs (Stumpf et al., 2008; Walsh et al., 2011; Zhong et al., 2019; Tang and Liu, 2020; Mirasbekov et al., 2021; Iwataki et al., 2022). Given the substantial climate-driven shifts in coastal ecological systems, accelerated interdisciplinary efforts are required to understand how climate change will influence the HABs.

This work applied a combination of satellite analysis and NGS to analyze the temperature trends of surface ocean water and reveal the presence of potentially toxic algae in the area of the marine benthic death (September-October 2020) reported near Avacha Bay Kamchatka region. We hypothesized that increased water temperature trend was the principal force behind algal blooms of potentially toxic algae in this area. The removal of seasonality and temperature trend projection suggest a 1.2-1.5^°^C temperature increase in critical for HABs development spring and summer months that favors further expansion of toxic algae to a north.

## Methods

### Sample collection

The surface seawater samples were collected a few days after the algal bloom in Avacha Bay, Kamchatka, and transported to the Nazarbayev University laboratory for further studies. The algal bloom event registered on 2020/06/26 followed by the mass death of marine life in September-October 2020 (**Fig. 1A, B**) (HAEDAT, 2021, reported by Orlova), and both were found to be spatially in line with the earlier recorded algal bloom event (Konovalova, and Syzykh, 2020).

**Figure 1.**
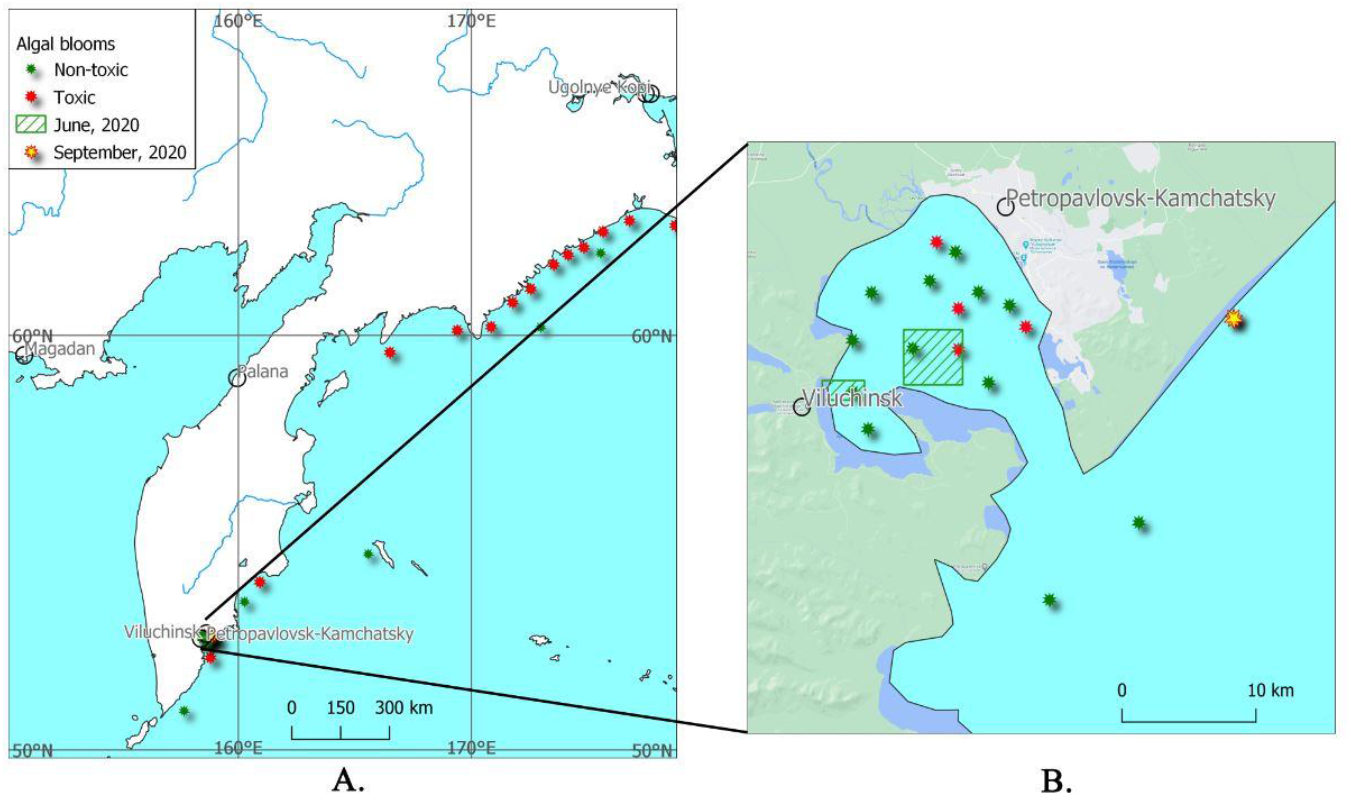
The observed algal blooms in Kamchatka region. **A**. The algal blooms in the Kamchatka region (data from Konovalova, 1993 combined with 2020 data events); **B**. The algal blooms at Avacha Bay in 2020, combined with data from Konovalova, 1993.

### Satellite analysis

The NASA/JPL product (2015) was considered during this study for monitoring the changes in the sea surface temperature. The Multi-scale Ultra-high Resolution (MUR) sea surface temperature (SST) analysis data were retrieved at 1 km x 1km grid (Chin et al., 2017). The dataset is the fusion of the AVHRR GAC, microwave, and in-situ sea surface temperature (SST) data. It consisted of the daily SST in Kelvin, which was then converted into Celsius degrees for this research using the following equation (1).

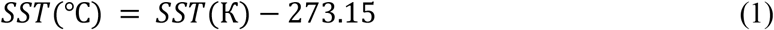

The derived data were used to plot the time series of the maximum SST. After a search in the USGS archive dataset, the cloudless Sentinel-2 image acquired on 2020/06/26 was used to calculate indices of the possible algal blooms and to monitor their spatial extents (Rawat et al., 2015). The Sentinel-2 are the optical ERS satellites launched and operated by the European Space Agency (ESA), covering the range of the spectral bands (**Table 1**).

**Table 1.** The characteristics of the Sentinel-2 bands (adapted from Wang et al., 2016)

| **Band number** | 1 | 2 | 3 | 4 | 5 | 6 | 7 | 8 | 8a | 9 | 10 | 11 | 12 |
| --- | --- | --- | --- | --- | --- | --- | --- | --- | --- | --- | --- | --- | --- |
| **Center (nm)** | 443 | 490 | 560 | 665 | 705 | 740 | 783 | 842 | 865 | 940 | 1375 | 1610 | 2190 |
| **Width (nm)** | 20 | 65 | 35 | 30 | 15 | 15 | 20 | 115 | 20 | 20 | 30 | 90 | 180 |
| **Spatial resolution (m)** | 60 | 10 | 10 | 10 | 20 | 20 | 20 | 10 | 20 | 60 | 60 | 20 | 20 |

According to the recent papers, the Maximum Chlorophyll Index (MCI) from the Medium Resolution Imaging Spectrometer (MERIS) data was used to detect surface and near-surface vegetation in the coastal and ocean water zones (Ansper and Alikas, 2019). After the MERIS mission was ended in 2012, nowadays, the Sentinel-2 is often being considered as an alternative source of environmental data, including the water surfaces. The MCI was calculated according to equation (2).

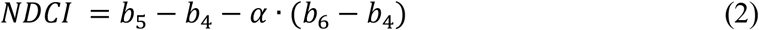

where the coefficient *α* was considered as the ratio of the difference of values for the bands 6, 5, and 4 (3).

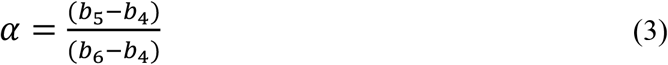

The Normalized Difference Chlorophyll Index was calculated as well for comparison purposes according to equation (4).

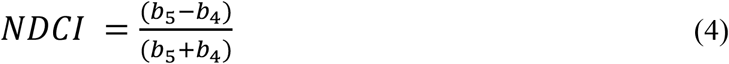

The algal bloom events reported earlier by other authors were mapped, and the algal bloom events possibly recorded in June and September 2020 were overlayed.

### DNA extraction

The PowerWater DNA Isolation Kit (Qiagen, USA) was used for DNA extraction from 1L of a water sample. Based on measurements of the Nanodrop spectrophotometer (ThermoFisher Scientific, USA) extracted DNA concentrations from “Vilyuchinskaya Bay” and “Avacha Bay” sampling sites (Location: **VB**: x 158.4408; y 52.90458; **AB**: x 158.8638; y 52.99816) were 20 ng/µl and 14 ng/µl, respectively.

### 16S rRNA gene sequencing

The methodology was adapted from the manufacturer’s instructions from Oxford Nanopore Technologies (further referred to as ONT) community. 16S Barcoding Kit (ONT, United Kingdom) was used for library preparation, which included PCR reactions and attachment of sequencing adapters. The reaction mixture included 10 ng of DNA sample, DreamTaq Hot Start PCR Master Mix (25 µl, Thermo Fisher Scientific, USA), 10 µM barcode primers (BC01-BC12) for 16S gene (1 µl, ONT, United Kingdom), and nuclease-free water (to reach 50 µL volume). There were two different samples from the Kamchatka region, namely “Vilyuchinskaya Bay” and “Avacha Bay,” and two different barcode primers were used to separate the results of the experiment. The reaction was set for initial denaturation (95 °C for 1 min); followed by 25 cycles of denaturation (95 °C for 20 sec), annealing (55 °C for 30 sec), extension (65 °C for 2 min); and ended by a final extension of 5 min at 65 °C. PCR products were cleaned up by AMPure XP beads (Beckman Coulter, USA), and PCR amplicons (with different barcodes) were mixed in equivalent amounts. Then, sequencing adapters were added to this mixture and incubated for 5 min at room temperature. The final DNA library also included loading beads and sequencing buffer (provided by the manufacturer). The MinION device (ONT, United Kingdom) was used for sequencing with flow cell R9.4. DNA library was transferred to the flow cell according to the manufacturer’s recommendations and the sequencing run continued for 12-24 hours. The base-calling of raw sequencing data was performed simultaneously during the run-in MinION sequencing device.

### 18S rRNA gene sequencing

The protocol was modified from a combination of a four-primer PCR procedure from the ONT community and the work of Hatfield and colleagues (2020). Most of the reagents were used from PCR-cDNA Barcoding Kit (ONT, United Kingdom). The amplification of the 18S rRNA gene was performed in two steps: the first reaction used a pair of tailed 18S primers for a small subunit ribosomal RNA gene; the second one was performed by barcoded universal primers with rapid attachment chemistry (supplied by the manufacturer).

The former PCR reaction mixture contained 5X Phusion HF buffer (10 μL, NEB, USA), 10 mM dNTPs (1 μL, Promega, USA), 10 μM each tailed 18S primers (2.5 μL, designed by Hatfield et al. (2020) and synthesized by Evrogen company, RF), Phusion HF DNA Polymerase (1 μL; NEB, USA), 100 ng of extracted DNA (depends on concentration) and nuclease-free water (to fill up to 50 μL). The reaction had the following steps (Hatfield et al., 2020): initial denaturation (98 °C for 1 min); 30 cycles of denaturation (98 °C for 10 sec), annealing (63 °C for 20 sec), extension (72 °C for 1.5 min); a final extension of 10 min at 72 °C. The presence of PCR-products with the expected size (about 3kb) was checked by gel electrophoresis stained by SYBR Safe DNA gel stain (ThermoFisher Scientific, USA).

Before the second reaction, PCR products were cleaned by Agencourt AMPure XP (Beckman Coulter, USA) with an elution volume of 10 μL (by 10 mM Tris-HCl pH 8.0 with 50 mM NaCl). All resulting products (10 μL) were used for further workflow. Based on the protocol from the ONT community, different barcoded primers (BP01-BP12, provided by ONT) were added to separate samples in the amount of 1 μL. Then, the LongAmp Taq 2X Master Mix (25 μL, NEB, USA) and nuclease-free water (14 μL) were added to reach a final reaction volume of 50 μL. The second reaction had the following steps: initial denaturation (94 °C for 2 min); 15 cycles of denaturation (94 °C for 15 sec), annealing (62 °C for 15 sec), extension (65 °C for 2.5 min); a final extension of 10 min at 65 °C. Further workflow of DNA library preparation and MinION sequencing was previously described in 16S rRNA gene sequencing above. The sequencing run continued for more than 20 hours. The base-calling was turned off during the run, and it was performed separately after the run.

### Bioinformatic analysis

The sequencing data was retrieved from the MinION device in archived fastq files (BioProject: PRJNA815633 Kamchatka_samples_2020; BioSamples:1. SAMN26631094: Vilyuchinskaya Bay; 2. SAMN26631097: Avacha Bay).

These sequences went through a quality check of a Phred quality score (further referred to as Qscore) of 7.0 by NanoFilt vs. 2.8.0 (De Coster et al., 2018). Then, qcat vs.1.1.0 (2021) was used to demultiplex sequencing reads and trim adapter sequences. The classification was performed in the Galaxy web platform using Kraken2 vs. 2.1.1 (Wood and Salzberg, 2014). Taxonomic labels were assigned to sequenced reads based on SILVA ribosomal RNA gene database (retrieved November 24, 2020). The Pavian vs.1.0 metagenomics data explorer (Brietweiser, and Salzberg, 2020) was used to visualize the results of the classification and to download taxonomic distributions for further analysis.

### Biodiversity indices

The biodiversity indices were used to assess relative taxa distribution within classification results, which enables a comparison of diversity between sampling sites. The first analysis included Shannon’s diversity index, which includes metrics based on both richness and dominance. Only dominance-based metrics of Simpson’s probability were applied in the study. Particularly, its inverse was calculated to have a positive correlation with Shannon Index. Both indices of diversity increase when the number of species or evenness increases. Only genus-level distribution was used to determine biodiversity.

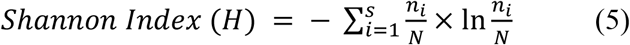

The equation (5) describes Shannon Index for distribution of taxa, where *s* is the total number of taxa identified after classification, *N* represents total number of sequencing reads and *n*_*i*_ indicates number of sequencing reads assigned to the taxon.

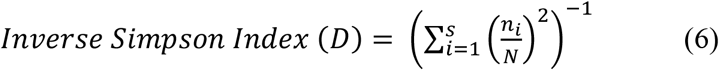

The equation (6) for Inverse Simpson Index for distribution of taxa, where *s* is the total number of taxa identified after classification, *N* represents total number of sequencing reads and *n*_*i*_ indicates a number of sequencing reads assigned to the taxon.

### Statistical analysis

For initial analysis of surface ocean temperature time series we applied the ordinary least square regression technique. Furthermore, due to the presence of significant seasonal component we used non-parametric algorithms for decomposition of time series with a significant seasonal component (Kovaldji et al., 2012).

Shortly, time series decomposition is a traditional approach presented in the form of equation:

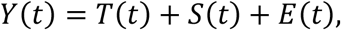

where *T(t)* is non-parametric trend, *S*(*t*)is non-parametric seasonal component, and *E*(*t*)is a noise. Two algorithms were used for decomposition of time series. Firstly, we used the smoothing algorithm based on local lineal regression (Kovaldji et al., 2012). Each point *t* was weighed using exponential coefficients calculated according the formula,

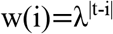

and WLS (weighted least squares) method was used for minimization of the values:

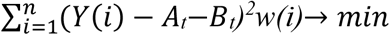

and calculation *Z(i)=A*_*t*_ *i + B*_*t*_ for each point.

Next, we introduced the curvature degree according to the formula:

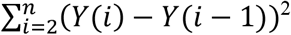

On the next step, time series (TS) was considered as a sum of trend and seasonal function. To find seasonal coefficients we minimized the degree of curvature for time series using the next equation for seasonal coefficients k(i):

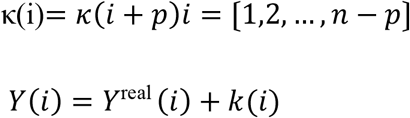

## Results

### 3.1 Satellite analysis

In our study, satellite data were used to detect sea surface water temperature (SST) changes during the summer of 2020 in the area near the Kamchatka coast (Avacha Bay). They were compared with satellite data of 2002-2020. The maximum sea surface temperature trends were derived from the SST product for 2002-2020 (**Suppl. Fig. 1A**) for 6705 days in total – a period from 2002/06/01 to 2020/10/09 (**Suppl. Table 1**). The time series (2002-2020) of the maximum SST values for April - September (**Suppl. Fig.1C-H**) were plotted. The maximum SST for 2020 was also plotted due to the events of the mass death of marine life in September 2020 (**Suppl. Fig.1B**). Further time series decomposition allowed to remove seasonal components (**Fig.2 A-F, Suppl. Data 1**) and to identify a stable increasing temperature trend in 2002-2020 period for May and July – 0.07^°^C and 0.05^°^C per year (**Fig. 2 C, E**). In March a stable trend for increase in ocean surface temperature started from 2011– for 0.08^°^ per year (**Fig.2 A**). The June and July 2020 peaks which were considered as anomalies affected the choice of the satellite images to monitor the chlorophyll indices.

**Figure 2.**
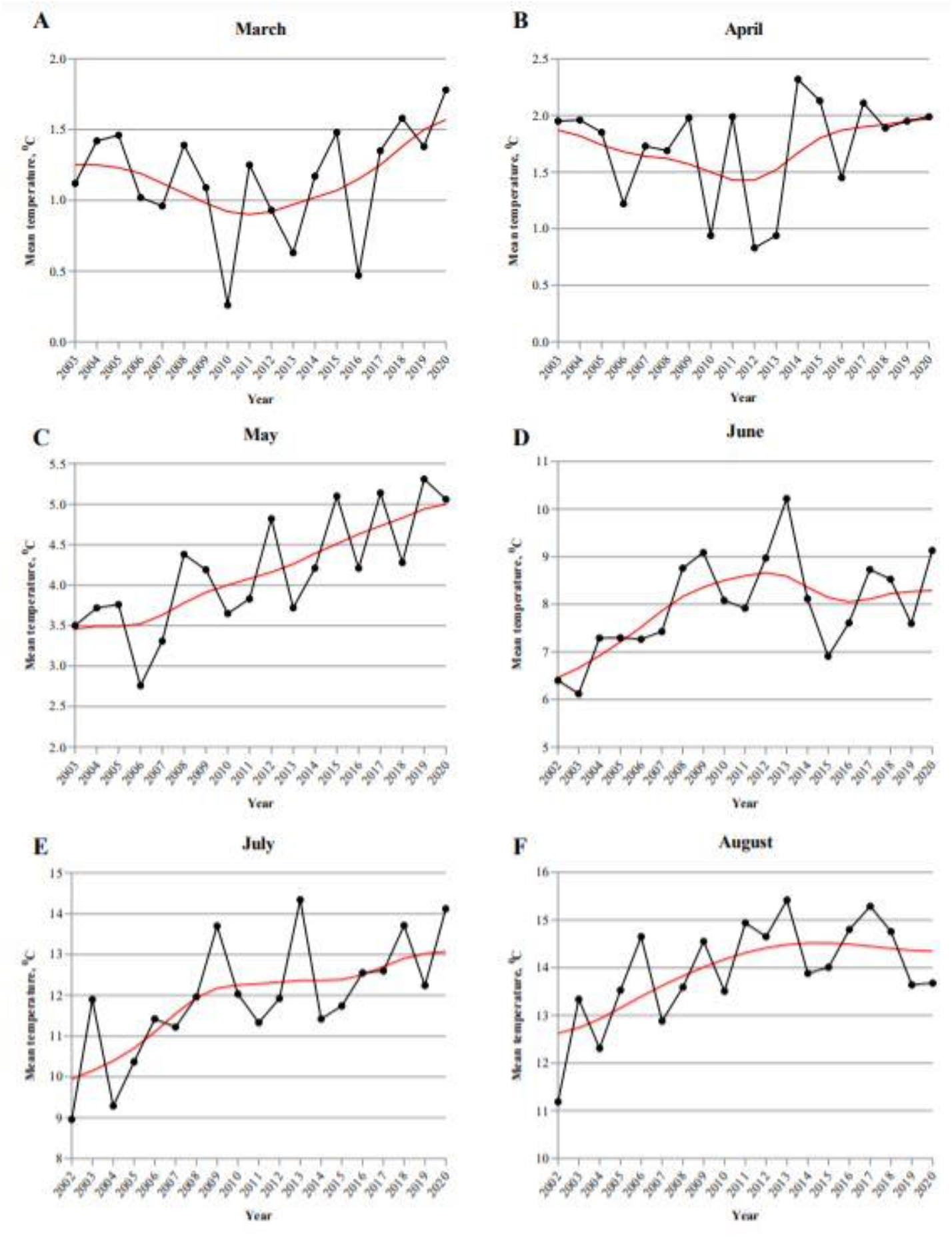
The ocean surface temperature (MUR-JPL-L4-GLOB-v4.1) at the Avacha Bay, Kamchatka (black line-initial time series; red line – seasonality free time series). **A**. March 2002-2020; **B**. April 2002-2020; **C**. May 2002-2020; **D**. June 2002-2020; **E**. July 2002-2020; **F**. August 2002-2020.

As the peaks of the maximum SST values for 2020 were in June (14.1 °C, 2020/06/30) and July (16.06 °C, 2020/07/28), the chlorophyll indices were calculated. The SST map was plotted for the anomaly temperatures in June 2020 (**Fig. 3B**). The date of the SST anomaly (2020/06/26) was nearly the acquisition date of the Sentinel-2 image (**Fig. 3B**). The sea surface temperature for Avacha Bay exceeded 13^°^C. The created true-color composition showed the structures on the sea surface attributed to the algal blooms (**Fig.3B**). The MCI with its maximum value of 834.9 (**Fig.3C**) and the NDCI with a maximum of 0.42 (**Fig.3D**) corresponded to those water surface structures and confirmed the detection of chlorophyll in those areas in the central and western part of Avacha Bay.

**Figure 3.**
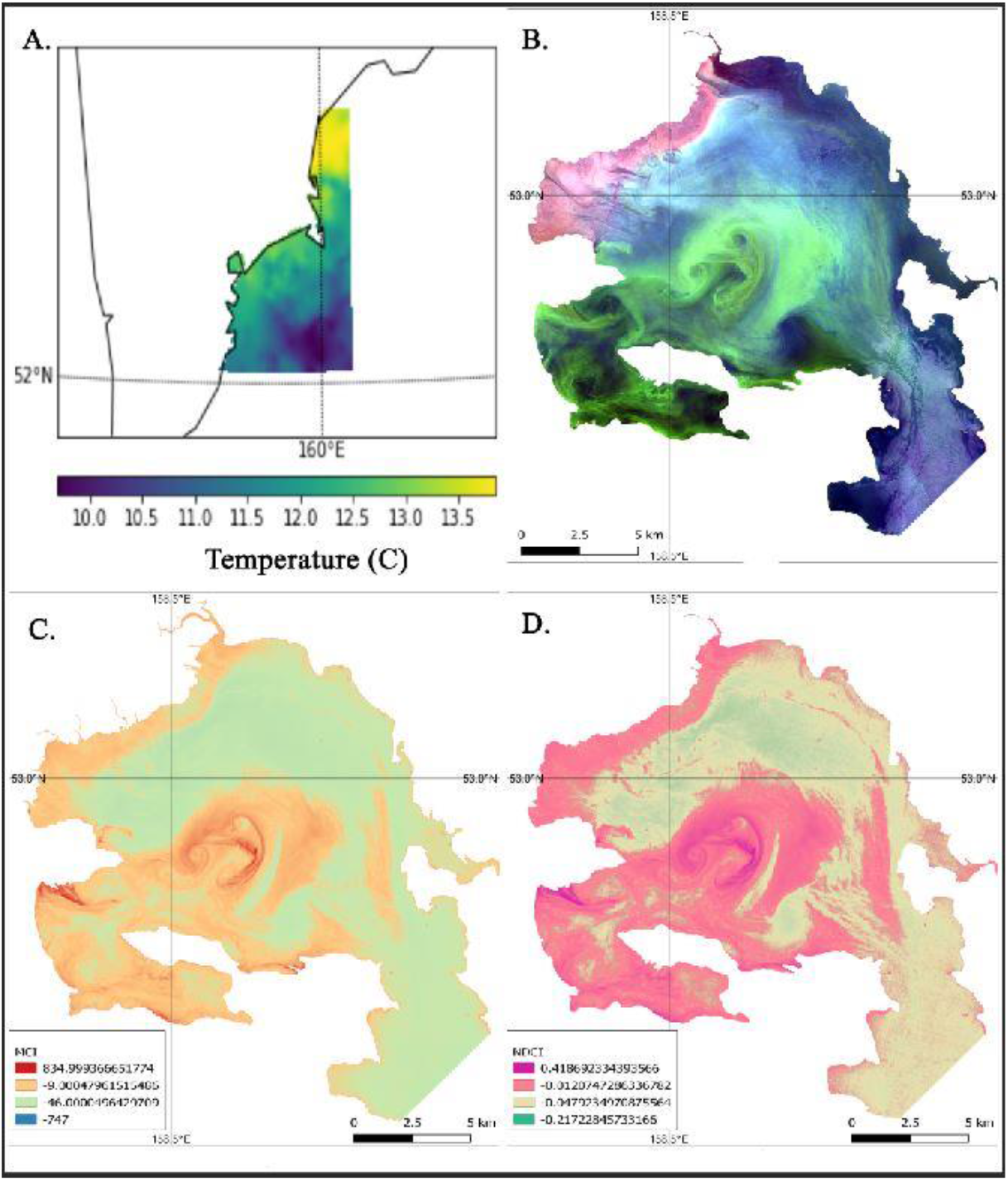
The properties of Avacha Bay observed from the space: **A**. The Sea surface maximum temperature 2020/06/28 (MUR-JPL-L4-GLOB-vs.4.1); **B**. The true-color image from the Sentinel-2, 2020/06/26; **C**. The Maximum Chlorophyll Index from the Sentinel-2, 2020/06/26; **D**. The Normalized Difference Chlorophyll Index from the Sentinel-2, 2020/06/26.

These facts could be evidence of the HAB processes’ intensification and need a detailed examination including time series of field observations.

### 3.2 Nanopore sequencing analysis

#### 3.2.1 16S rRNA gene sequencing

The sequencing run was repeated twice to have an equivalent amount of sequencing information of 16S amplicons from both sampling sites. After quality checks (Qscore > 7), demultiplexing and trimming of adapter sequence, a number of sequencing reads were 388,162 and 401,287 based on assigned barcodes for “Vilyuchinskaya Bay” and “Avacha Bay”, respectively. These datasets (data BioProject: PRJNA815633 available at https://dataview.ncbi.nlm.nih.gov/object/PRJNA815633?reviewer=qmvrh3095np80tn7g9t1oqj30g) were used for the taxonomic analysis of samples based on the SILVA SSU database, which resulted in more than 99% classification of reads. Details of sequencing data are summarized in **Suppl. Table 2**. Distributions at different taxonomic levels are illustrated in **Figure 4A–B**. According to this data, both of the samples had a high percentage of sequencing reads from Proteobacteria (65-80%) and Bacteriodota (11-16%) phyla. The “Avacha Bay” sample site had a higher proportion of Cyanobacteria and Verrucomicrobiota taxa compared to the “Vilyuchinskaya Bay” sample site. Other phyla represent less than 1.5% of data from both sequencing runs. The significant difference was the prevalence of Campylobacterota phylum (20.7%) in the “Vilyuchinskaya Bay” sample when the “Avacha Bay” sample indicated this phylum in the smaller proportion (about 0.05% of classification results). At the genus level, 1,893 different genera were identified from two sampling sites, 998 of which are present in both of the samples. **Figure 4A– B** shows ten genera with the highest number of reads, which represents around 65% of whole sequencing data. Among these selected results, only *Flavobacterium* genus was found in both of the sampling sites, and the percentage distribution of other genera was entirely different between samples.

**Figure 4.**
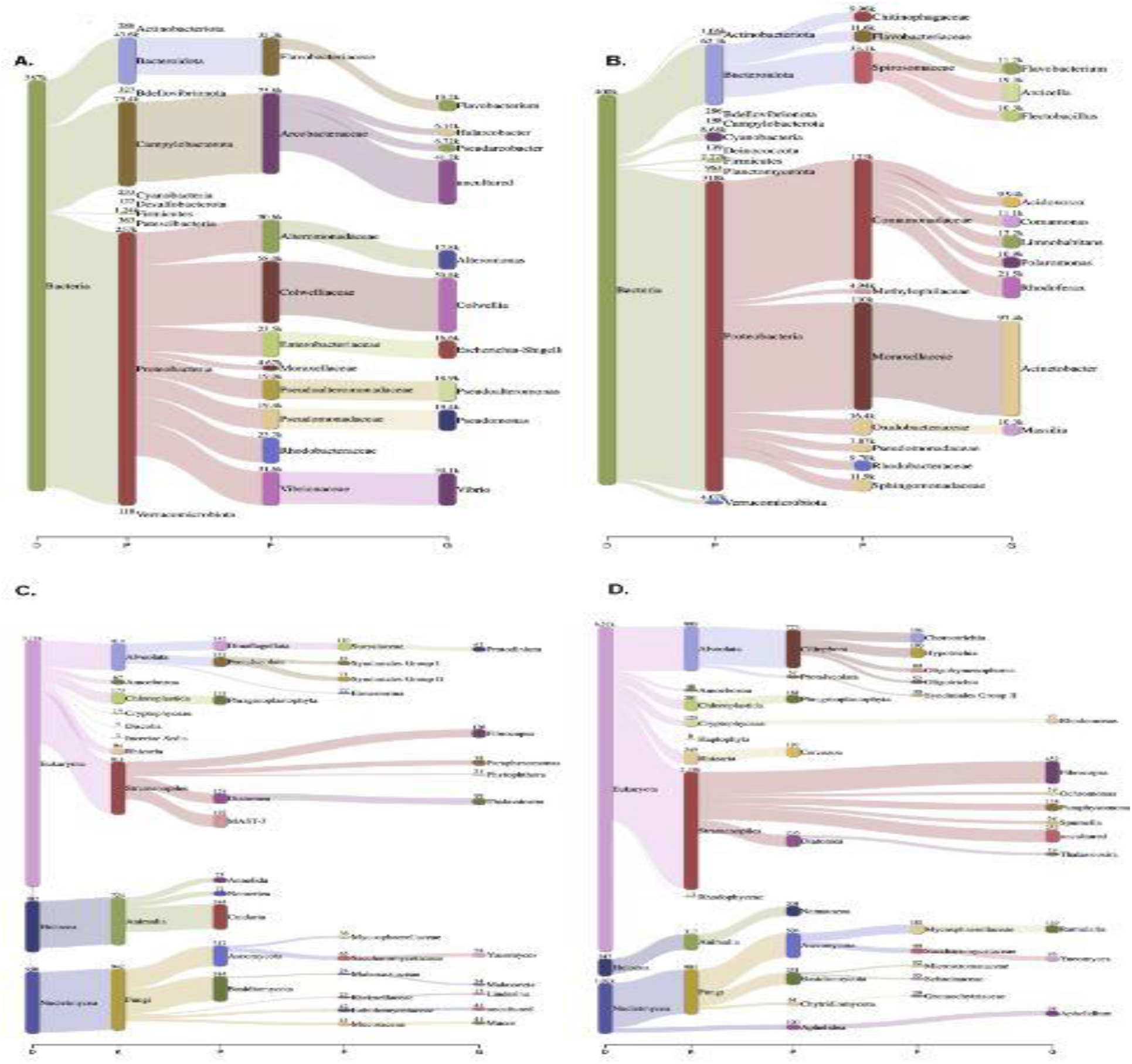
Phylogenetic distribution of Kamchatka samples based on results of nanopore sequencing of two different target genes: 16S rRNA gene (**A** – **B**) and 18S rRNA (**C – D**). Samples were taken from “Vilyuchinskaya Bay” (**A** and **C**) and “Avacha Bay” sampling sites (**B** and **D**). **A**. Sankey diagram of kraken2 report for taxonomic assignment based on SILVA database. Sequenced reads are represented at different phylogenetic levels: domain (D), kingdom (K), phylum (P), family (F) and genus (G) (top 10 taxa at each level). Amount of sequencing reads from nanopore sequencing is indicated in numbers, which is proportional to the width of the flow.

To compare biodiversity between these two samples, the Shannon’s diversity index (*H*) and Simpson index (*D*) was performed at a 97% cut-off. The cut-off was applied for these analysis methods to remove taxa with a low number of reads. According to the results, the bacterial biodiversity index in “Vilyuchinskaya Bay” (*H* = 3.489; *D* = 14.661) was bigger than in “Avacha Bay” (*H* = 3.433; *D* = 10.001) based on 16S rRNA gene sequencing.

#### 3.2.2 18S rRNA gene sequencing

After the sequencing run, raw sequencing data went through the same processing pipeline mentioned in the previous section, and the number of sequencing reads was 198,645 and 105,152 for “Vilyuchinskaya Bay” and “Avacha Bay,” respectively. The majority of these reads had a small read length, but the amplicon size was estimated to be about 3kb (Hatfield et al., 2020). Therefore, an additional read length filter was applied to leave sequences with the length of 500 base pairs and longer. It resulted in 8,699, and 9,624 sequencing reads for samples from “Vilyuchinskaya Bay” and “Avacha Bay,” respectively. The data was used for classification based on the SILVA SSU database, which resulted in 40-70% classification. Summary of sequencing data analysis is found in **Suppl. Table 2**. Distributions at different taxonomic levels are illustrated in **Figure 4C– D**. When comparing ten genera with the highest proportion of sequencing reads in “Vilyuchinskaya Bay” and “Avacha Bay,” between microalgae, there are three intersecting genera: *Fibrocapsa, Thalassiosira*, and heterotrophic *Paraphysomonas*; and the former being the most abundant in both sampling sites. Other most abundant genera are illustrated in **Figure 4C– D**. Altogether these listed genera represent around 40-50% of sequencing data. The biodiversity index in “Vilyuchinskaya Bay” (H=4.698; D=40.369) was bigger than in “Avacha Bay” (H=4.358; D=23.458) based on the eukaryotic classification of 18S rRNA sequencing results. Genera that include potentially toxic and a bloom-forming micro-algae were identified based on previous reports from the Kamchatka region (**Table 2**).

**Table 2.** Summary of intersecting genera with the list of harmful algae in the Far Eastern Seas of Russian Federation (from Orlova et al., 2010).

| Group name | Genus name | Number of seq. reads in “Vilyuchinskaya Bay” | Number of seq. reads in “Avacha Bay” |
| --- | --- | --- | --- |
| Raphidophytes | Fibrocapsa* | 126 | 452 |
| Diatoms | Thalassiosira† | 99 | 59 |
|  | Amphora* | 7 | 20 |
|  | Chaetoceros† | 2 | 6 |
|  | Rhizosolenia† | 2 | 3 |
|  | Skeletonema† | NA | 2 |
|  | Proboscia† | NA | 1 |
| Chrysophytes | Dinobryon† | 2 | 19 |
| Dinoflagellates | Gonyaulax* | 4 | 2 |
|  | Gymnodinium* | 1 | NA |
|  | Karlodinium* | 1 | 1 |
|  | Prorocentrum*† | NA | 1 |
|  | Protoperidinium*† | NA | 1 |
|  | Amphidinium* | NA | 1 |

\* Genera that contain potentially toxic species.
† Genera that contain bloom-forming species.

## Discussion

Global ocean temperature has already increased and will continue to warm during the 21st century (IPCC, 2013). Surface waters warmed extremely rapidly in some coastal areas, already reaching conditions projected in year-2100 (Pershing et al., 2021) and inducing corresponding shifts in all ecosystem levels (Record et al., 2021). Temperature is a major driver of phytoplankton biogeographic boundaries (Thomas et al., 2012; Litchman et al., 2015), and its increase is associated with the increase of HABs duration and range (Moore et al., 2008; Hallegraeff, 2010; Ralston et al., 2014).

Based on satellite data analysis from Sentinel-2, we demonstrated a significant change in the surface water temperature in the Kamchatka region near Avacha Bay in 2002-2020, in the spring and summer months (in particular, May and July) (**Fig.2C, E**). Projected in the next 15 years it will bring increase in the surface temperature 1.2-1.5^°^C. It is in line with the research of Rogachev and Shlyk (2021), who analyzed historical CTD and Argo observations and found out that temperature in the Kamchatka Current core increased from 1990 up by 2.4^°^C from the beginning of observations. Furthermore, analysis of chlorophyll distribution in July 2020 based on Sentinel-2 data had confirmed an algal bloom in Avacha Bay (**Fig.3**). The Kamchatka peninsula has periodical HABs (Konovalova, 1993; Orlova et al., 2010), including a recent event near Avacha Bay in September-October 2020 that was reported to the Harmful Algae Event Database (2021; reported by Orlova).

The gradient in temperature growth responses favors the enhanced development of HABs at higher latitudes. In the study of Brandenburg and coauthors (2019), HAB species from higher latitudes responded more positively to high temperatures, while algal isolates from lower latitudes responded negatively. This is in line with the earlier observation that species from higher latitudes grow below their temperature optimum while species from tropics grow at their temperature optimum (Thomas et al., 2012; Boyd et al., 2013). One of the examples of the increasing temperature gradient effect for potentially toxic algae distribution is an expansion in the last two decades of HABs of *Karenia mikimotoi* along the coast of China (longitudinal range of 200) (Baohong et al., 2021). Another example is a temperature change in the surface ocean waters near the Gulf of Maine (Pershing et al., 2021) that led to the northward appearance of blooms of *K. mikimotoi* and diatom *Pseudo-nitzchia australis* (Record et al., 2021).

We assessed the 16S bacterioplankton and 18S microalgae plankton communities’ composition following the marine life kill event in September-October 2020 and identified a presence of practically all potentially toxic genera reported earlier to be responsible for HABs events near the Kamchatka peninsula (Orlova et al., 2010). The climate change and increase of temperature in this region can produce algal blooms originating from cysts coming from the Bering Sea (*Alexandrium* spp.) and from more temperate areas - *Karenia* spp. Earlier *Alexandrium* spp. were related to a number of PSPs events on the Russian side of the Bering strait and near the Kamchatka peninsula (Konovalova, 1993; Selina et al., 2006). Though HABs events mainly report one dominating phytoplankton species, with future development of detection methods, information surfacing about HABs where multiple algal species may be involved (Mulholland et al., 2014; Paerl et al., 2018), and co-occurrence of various toxins may happen (Peacock et al., 2018). Thus, during *Karenia* PSP-outbreaks in New Zealand, there was confusion as to how many *Karenia* species were present (Bates et al., 1993; Chang, 1995), and at least two *Karenia* species were reported -*K*. cf. *brevis* and *K*.*mikimotoi*. In the New Zealand outbreaks ten years later, in 2002 and 2003, *K*.cf. *brevis* bloomed together with *K*.*mikimotoi* and *K. brevisculata* (Chang, 2003). Moreover, toxic algae species may lead to a harmful event at different cell concentrations. Thus, toxic *Alexandrium* spp. leads to shellfish contamination even at very low concentrations such as 200 cells per liter - making it challenging to use satellite analysis for HABs prediction (Eckford-Sopper et al., 2013). In contrast, *Karenia* spp., a dinoflagellate that causes ‘red tide,” typically accumulates at high concentrations that discolor the water. There is no direct correlation between a species initial population density and its bloom-forming capacity, partly because even rare species can reach an astronomically high abundance-therefore, longitudinal time-series data are needed. It leaves unresolved questions regarding what processes select the species that proliferate during HABs (Paerl et al., 2018; Wells et al, 2020): If species are chosen by “precedent and stochasticity”(Reynolds et al., 2000), how will the change in temperature range and anthropogenic factors affect multi-species bloom development?

In addition to changes in environmental factors along the Kamchatka coast (storms, winds) that favors HABs (Laanaia et al., 2013; Kamiyama et al., 2014), there are physiological characteristics that support the expansion of *Karenia* blooms to the north. They include (1) capabilities to utilize anthropogenically derived inorganic and organic nitrogen and phosphorus compounds (Yamaguchi, and Itakara, 1999); (2) degradation of the marine environment directly associated with human activities such as coastal embayments, pollution, over-fishing and over-exploitation of natural resources (Glibert et al., 2005). The Avacha Bay (Kamchatka) reported having the most unfavorable situation in terms of pollution (Rostove, and Rudykh, 2018).

Recently, Kuroda and colleagues (2021) described large scale HABs in coastal waters of Hokkaido, Japan in mid-to-late September 2021. Moreover, Iwataki and co-authors (2022) used morphology and phylogenetic reconstruction based on the large sub-unit ribosomal nucleotide (LSU rDNA) and Internal Transcriber Spacer (ITS) to demonstrate that *Karenia selliformis* sequences from 2021 Hokkaido HABs were identical to the sequences from the marine benthic kill in Avacha Bay, Kamchatka, 2020. *Karenia selliformis* blooms have been reported in coastal waters of New Zealand, Mexico, Tunisia, Kuwait, Iran, China (Mardones et al., 2020), but first time in Japan and Kamchatka. It is known that *Karenia* sp. blooms are driven by anthropogenic interference and temperature increase (Feki et al., 2013). Thus, *K*.*selliformis* first time recordings such as in the Gulf area (Heil et al., 2001) was suspected alien species introduction related to ballast water problem (Al-Yamani et al., 2015; Clarke et al., 2019). Similarly, the first recording of *K*.*selliformis* occurrence in Japan and Kamchatka waters (Iwataki et al., 2022) may also be possible alien species introduction in Avacha Bay region. The International Convention for the Control and Management of Ship’s Ballast Water and Sediments (BWM Convention 2004) entered in force in 2017 but by no means provide a fully effective solution to problem of HABs related to ballast water (Hallegraeff, 2015).

We hypothesize that long-term temperature trends in Avacha Bay surface waters may lead to increase of HABs frequency of multiple toxic algae taxa. HAB species from high latitudes (*Alexandrium* sp.), *Karenia* sp. brought to the area from different locations and other potentially toxic algal taxa suggest a potential threat of recurrent blooms and risk of further spread of this species along Kamchatka and in Japanese coastal waters. More attention should be given to mixed HABs and analytical methods for their research such as eDNA monitoring. Number of nonindigenous species introduced in the area (with ballast water or otherwise) or colonization pressure may increase an invasion risk with new toxic algae species (Lockwood et al., 2009; Briski et al., 2014).

## Conclusions

We had demonstrated using Sentinel-2 satellite analysis an increasing temperature trend in surface water temperature in Avacha Bay of Kamchatka peninsula that may lead to an increase of frequency and intensity of HABs in the region. The September-October 2020 benthic marine life kill was preceded by an extreme heatwave and extensive algal bloom in July 2020. Nanopore-based next-generation sequencing with MinION platform revealed a presence of multiple potentially toxic and bloom-forming algae from genera overlapping with past HABs dominant species in the region of the Kamchatka peninsula. Further studies need to assess the presence of multiple toxic algal species in Avacha Bay, Kamchatka and to undertake regionally focusing climate-change related risk assessment. Temperature trend projection suggest with some certainty ocean surface water warming 1.2-1.5^°^C in the spring and summer months in the next decade and increase of HABs frequency in coastal regions of Kamchatka.

## Abbreviations

HABs: Harmful Algae Blooms
HAEDAT: Harmful Algae Event Database
MCI: Maximum Chlorophyll Index
MERIS: Medium Resolution Imaging Spectrometer
MUR-JPL-L4-GLOB-vs.4.1: Multiscale Ultrahigh Resolution Level 4 Global sea surface temperature dataset at the Jet Propulsion Laboratory (NASA) vs. 4.1
NEB: New England Biolaboratories
NGS: next-generation sequencing
NDCI: Normalized Difference Chlorophyll Index
ONT: Oxford Nanopore Technologies
PSP: paralytic shellfish poisoning
PCR: polymerase-chain reaction
SST: sea surface water temperature

## Acknowledgements

Funding for this work came from Nazarbayev University grant, FCDGRP #110119FD4513 to N.S.B. We are grateful to colleagues for sharing water samples from Avachin Bay area.

## Legends to Supplemental Data

**Supplemental figure 1**. The maximum Sea surface temperature (MUR-JPL-L4-GLOB-v4.1) at the Avacha Bay. Kamchatka. **A**. 2002-2020; **B**. 2020; **C**. April 2002-2020; **D**. May 2002-2020; **E**. June 2002-2020; **F**. July 2002-2020; **G**. August 2002-2020; **H**. September 2002-2020.

**Supplemental Data 1**. Temperature data and trends after pre-processing and seasonality removing.

